# Development of a Nested Association Mapping (NAM) population for untangling complex traits in lentil (*Lens culinaris* Medik.)

**DOI:** 10.1101/2025.06.17.660011

**Authors:** Sandesh Neupane, Robert Stonehouse, Larissa Ramsay, Mari Baker, Teketel Haile, David J. Konkin, Christine Sidebottom, Kirstin E. Bett

## Abstract

Understanding the genetic basis of complex traits remains a key challenge in crop improvement. This study aimed to develop a structured, multi-parental mapping population to enhance the resolution of quantitative trait dissection, based on the hypothesis that a Nested Association Mapping (NAM) design would enable the detection of minor-effect loci often overlooked by traditional biparental or diversity panel-based approaches. A lentil (*Lens culinaris* Medik.) NAM population was developed by crossing the Canadian cultivar CDC Redberry with 32 diverse genotypes sourced globally from three major lentil-growing macro-environments: Northern temperate, Mediterranean, and South Asia. The resulting recombinant inbred lines were phenotyped for key phenological traits, days to emergence (DTE), flowering (DTF), and maturity (DTM), under field conditions, and genotyped using exome capture sequencing. Genome-wide association studies for DTF identified 14 significant loci across six chromosomes, including the known *FTb* locus and novel associations near *AP3a, HUB2a*, and *PIF6* genes. These results demonstrate the utility of the NAM design in detecting both major and minor-effect loci that underlie complex trait variation. To our knowledge, this is the first publicly available NAM population in lentil. It provides a high-resolution, globally representative platform for trait discovery, pre-breeding, and collaborative genetic improvement of this nutritionally and agronomically important legume.

## 1. Introduction

Broadening our understanding of the genetic components of complex traits, known as quantitative genetics, is a fundamental pursuit in contemporary plant breeding and genetics. These traits, characterized by continuous variation and influenced by numerous genes and environmental factors, present substantial challenges to crop improvement efforts, especially when the goal is to broaden the genetic base of breeding populations (Holland, 2007). Traditional genetic mapping methods, including bi-parental quantitative trait loci (QTL) mapping and genome-wide association studies (GWAS), have played pivotal roles in identifying genomic regions linked to complex traits. However, these methods have inherent limitations; QTL mapping often offers limited resolution due to restricted recombination events, while GWAS can be confounded by population structure, potentially causing false associations and reducing the detection power for minor-effect loci (Flint-Garcia et al., 2003; Yu et al., 2008).

To overcome these limitations, multi-parental populations such as Multi-parent Advanced Generation Intercross (MAGIC) and Nested Association Mapping (NAM) have emerged as innovative strategies to enhance genetic resolution and increase the power of association studies (Mackay et al., 2014; Yu et al., 2008). NAM, initially developed for maize by Yu et al. (2008), combines the strengths of GWAS and QTL mapping while mitigating their respective drawbacks. The effectiveness of NAM has been demonstrated in several crops, including maize, wheat, sorghum, and peanut, significantly contributing to genetic discovery and breeding advancements (Bouchet et al., 2017; Gangurde et al., 2020; Kidane et al., 2019; Yu et al., 2008). However, the application of NAM populations remains relatively unexplored in minor yet important crops like lentil (*Lens culinaris* Medik.).

Lentil is a legume crop globally recognized for its high protein content, dietary fibre, vitamins, and minerals. Lentil also plays a vital role in sustainable agriculture by aiding in converting atmospheric nitrogen (N) to crop-available N through N-fixation. Despite its agricultural importance, lentil breeding programs often face challenges, particularly in introducing new genetic diversity to their production regions due to interactions between the growing environment and flowering time genes (Neupane et al., 2022; Summerfield et al., 1985; Wright et al., 2021). While recent lentil genetics studies have made strides in identifying genetic markers for key phenological traits, these studies have predominantly used either diversity panels or bi-parental populations, such as recombinant inbred lines (RILs) (Fedoruk et al., 2013; Fratini et al., 2007; Haile et al., 2021; Kahriman et al., 2015; Lake et al., 2024; Neupane et al., 2022; Sarker et al., 1999; Tullu et al., 2008) and typically lack the resolution required to pinpoint minor-effect loci.

In this context, the objective of this study is to describe the development of a NAM population based on a diverse set of lentils that was then genotyped and is useful for dissecting complex traits. It is hypothesized that the NAM population structure would enhance the detection of minor-effect genes and refine the genetic resolution of complex trait such as flowering time control in lentil.

## 2. Materials and Methods

### 2.1 Description of parental lines, population development and their field evaluation The lentil Nested Association Mapping (NAM) population was developed by crossing CDC

Redberry, a high-yielding, red cotyledon Canadian cultivar with a reference genome (Ramsay et al., 2021), with 32 diverse genotypes sourced from different lentil-growing regions worldwide (Table 1). These other genotypes were selected to represent the three major macro-environments for lentil cultivation: Northern temperate, Mediterranean, and South Asia (Khazaei et al., 2016; Wright et al., 2021). Each cross of CDC Redberry with 32 other genotypes resulted in F1 plants which were selfed to produce the F2 generation, initiating the development of recombinant inbred lines (RILs) through single seed descent (Table S1, Figure S1).

**Table 1.** Summary of the lentil NAM population, including pedigree information, origin of the second parent and the total number of lines developed and sequenced.

| Sub-Population Name | Pedigree | Second Parent Origin | # RILs |
| --- | --- | --- | --- |
| NAM-1 | CDC Redberry x 2861-15a | Canada | 127 |
| NAM-2 | CDC Redberry x 3339-3 | Canada | 119 |
| NAM-3 | CDC Redberry x 2275-15 | Canada | 128 |
| NAM-4 | CDC Redberry x Shasta | USA | 115 |
| NAM-5 | CDC Redberry x 02M-12 | Canada | 54 |
| NAM-6 | CDC Redberry x 3954-105 | Canada | 51 |
| NAM-7 | CDC Redberry x Guarena | Spain | 54 |
| NAM-8 | CDC Redberry x 5365-6 | Canada | 50 |
| NAM-9 | CDC Redberry x 4656-3 | Canada | 58 |
| NAM-10 | CDC Redberry x Lupa | Spain | 61 |
| NAM-11 | CDC Redberry x ILL 213 | Afghanistan | 59 |
| NAM-12 | CDC Redberry x ILL 1706 | Ethiopia | 56 |
| NAM-14 | CDC Redberry x ILL 5722 | Syria | 59 |
| NAM-15 | CDC Redberry x ILL 6002 | Syria | 58 |
| NAM-16 | CDC Redberry x ILL 7727 | Morocco | 56 |
| NAM-17 | CDC Redberry x ILL 1704-1 | Ethiopia | 56 |
| NAM-18 | CDC Redberry x LR59-81 | Canada | 55 |
| NAM-20 | CDC Redberry x Ranjan | India | 60 |
| NAM-21 | CDC Redberry x ILL 8007 | Bangladesh | 54 |
| NAM-22 | CDC Redberry x ILL 1959 | Ethiopia | 55 |
| NAM-24 | CDC Redberry x IG 858 | Algeria | 58 |
| NAM-25 | CDC Redberry x ILL 11551 | India | 58 |
| NAM-26 | CDC Redberry x ILL 4609 | Pakistan | 59 |
| NAM-27 | CDC Redberry x ILL 618 | Tajikistan | 53 |
| NAM-28 | CDC Redberry x ILL 9 | Jordan | 60 |
| NAM-29 | CDC Redberry x ILL 927 | Turkey | 60 |
| NAM-30 | CDC Redberry x PI 182217 | Turkey | 60 |
| NAM-31 | CDC Redberry x PI 426797 | Pakistan | 60 |
| NAM-32 | CDC Redberry x PI 429838 | Iran | 54 |
| NAM-33 | CDC Redberry x PI 431884 | Iran | 55 |
| NAM-34 | CDC Redberry x PI 432005 | Iran | 58 |
| NAM-35 | CDC Redberry x PI 432245 | Lebanon | 59 |

The entire population of 2,079 lines including parents was then evaluated under field conditions at Sutherland, SK (52.17° N, 106.52° W) during the summer of 2019. The field experiment was conducted using an augmented design with a single 1 m row for each genotype. Days to emergence (DTE) - when 10% of plants in a row had emerged from the soil surface, days to flowering (DTF) - when 10% of plants in a row had at least one open flower, and days to maturity (DTM) - when 10% of plants in each row displayed 50% pod maturity, were recorded for each line.

### 2.2 Genotyping and sequencing

Leaf tissue was collected for DNA extraction from the last single plant before bulking seed to create a final RIL. Genotyping was conducted using a lentil exome capture assay (Ogutcen et al., 2018) followed by skim sequencing.

DNA Libraries were prepared from the parents and lines of the NAM population using the [Illumina](https://support.illumina.com/sequencing/sequencing_kits/illumina-dna-prep/documentation.html) Nextera^TM^ DNA Flex protocol. Whenever given an option, shaking was the method chosen to mix the samples. During the Post Tagmentation Cleanup, we followed the steps recommended for processing more than forty-eight samples simultaneously. For the post-ligation PCR Program, we used nine cycles of amplification. The target fragment size was 350 base pairs. The quality and yield of the libraries were checked on an Agilent Technology 2100 Bioanalyzer using a High Sensitivity DNA chip kit.

Exome captures were performed using the SeqCap EZ HyperCap Workflow v2.1 (Roche) with minor modifications. Equal volumes of 96 Nextera^TM^ DNA Flex libraries were combined such that the total amount of DNA pooled equaled 1 µg. Half of the recommended amount of probe was added to the hybridization buffer. Fourteen amplification cycles were used in the post-capture PCR. To purify the post-capture amplified DNA, the libraries were bound to 17.5 µl of AMPure XP beads. The quality of the Post-Capture Library pools was checked on an Agilent Technology 2100 Bioanalyzer using a High Sensitivity DNA chip kit. Libraries were quantified on a Qubit^TM^ (Invitrogen^TM^). Exome capture library pools were evenly divided and sequenced on 3 lanes of a NovaSeq 6000 (Illumina; S4 flowcells, 2×150 bp reads) which resulted in approximately 1.0X coverage for each NAM line.

Alignment and variant calling were performed using the previously established exome capture processing workflow (Ogutcen et al., 2018) using the CDC Redberry lentil genome (Lcu.2RBY; Ramsay et al., 2021) as a reference. SNPs with greater than 35% missing data, minor allele frequencies below 5%, or a PHRED quality score below 30 were excluded.

### 2.3 Genome-wide association studies for flowering time

To evaluate the power of the NAM population in identifying candidate genes or markers associated with complex traits, a genome-wide association study (GWAS) was performed on the entire NAM population. The study used genotypic data and phenotypic observations of days to flowering (DTF), collected from a field trial conducted in Sutherland, Saskatoon, SK, in 2019.

GWAS was implemented using the Bayesian-information and Linkage-disequilibrium Iteratively Nested Keyway (BLINK) model, which provides greater statistical power and computational efficiency over other single and multi-locus statistical models (Huang et al., 2019; Wang & Zhang, 2021). Analyses were executed using the Genome Association and Prediction Integrated Tool (GAPIT, version 3.0) (Wang & Zhang, 2021) in R (RCoreTeam, 2019). A stringent significance threshold of −log10(0.05/ number of markers used), derived from a Bonferroni correction (Holm, 1979) for multiple testing across all markers, was applied to rigorously control the false discovery rate.

## 3. Results and Discussion

### 3.1 Establishment of lentil NAM population

A lentil Nested Association Mapping (NAM) population was developed by crossing the elite Canadian cultivar CDC Redberry with 32 diverse accessions representing major lentil-producing macro-environments: Northern temperate, Mediterranean, and South Asia (Table 1). This whole process yielded 2,079 lines with 50 to129 RILs per cross combination. This multi-parental NAM population expands the genetic base available to lentil breeding programs. NAM designs increase recombination and allele diversity compared to traditional biparental populations, while population structure is controlled via the common parent (CDC Redberry, in our case) (Yu et al., 2008; Mackay et al., 2014). Such designs have proven effective in dissecting complex traits and utilizing underrepresented germplasm in other crops, including maize, wheat, and soybean (Bouchet et al., 2017; Song et al., 2017; Kidane et al., 2019).

### 3.2 NAM reflected similar phenotypic variation as diversity panel

Field evaluation of the NAM population revealed substantial phenotypic variation in key phenological traits, days to emergence (DTE), days to flowering (DTF), and days to maturity (DTM) (Figure 1). As previous studies have highlighted, DTF is a key phenological trait influenced by photoperiod and temperature, which are crucial environmental parameters for determining adaptation to specific agro-climatic zones (Wright et al., 2021), a deeper analysis was conducted on DTF. A marked variation was observed both within and among RIL families when compared to the common parent (CDC Redberry) and their respective donor parent. Some families displayed broad ranges in DTF, while others showed narrower distributions. Notably, RIL families with a second parent from temperate regions tended to flower later, whereas those derived from South Asian second parents flowered earlier. Families with Mediterranean-origin second parents exhibited a mixture of early- and late-flowering genotypes. These broad phenotypic variations across diverse genetic backgrounds confirm that the NAM population captures substantial genetic diversity. This observation aligns with results from the lentil diversity panel, where similar patterns of variation in DTF were reported (Wright et al., 2021).

**Figure 1.**
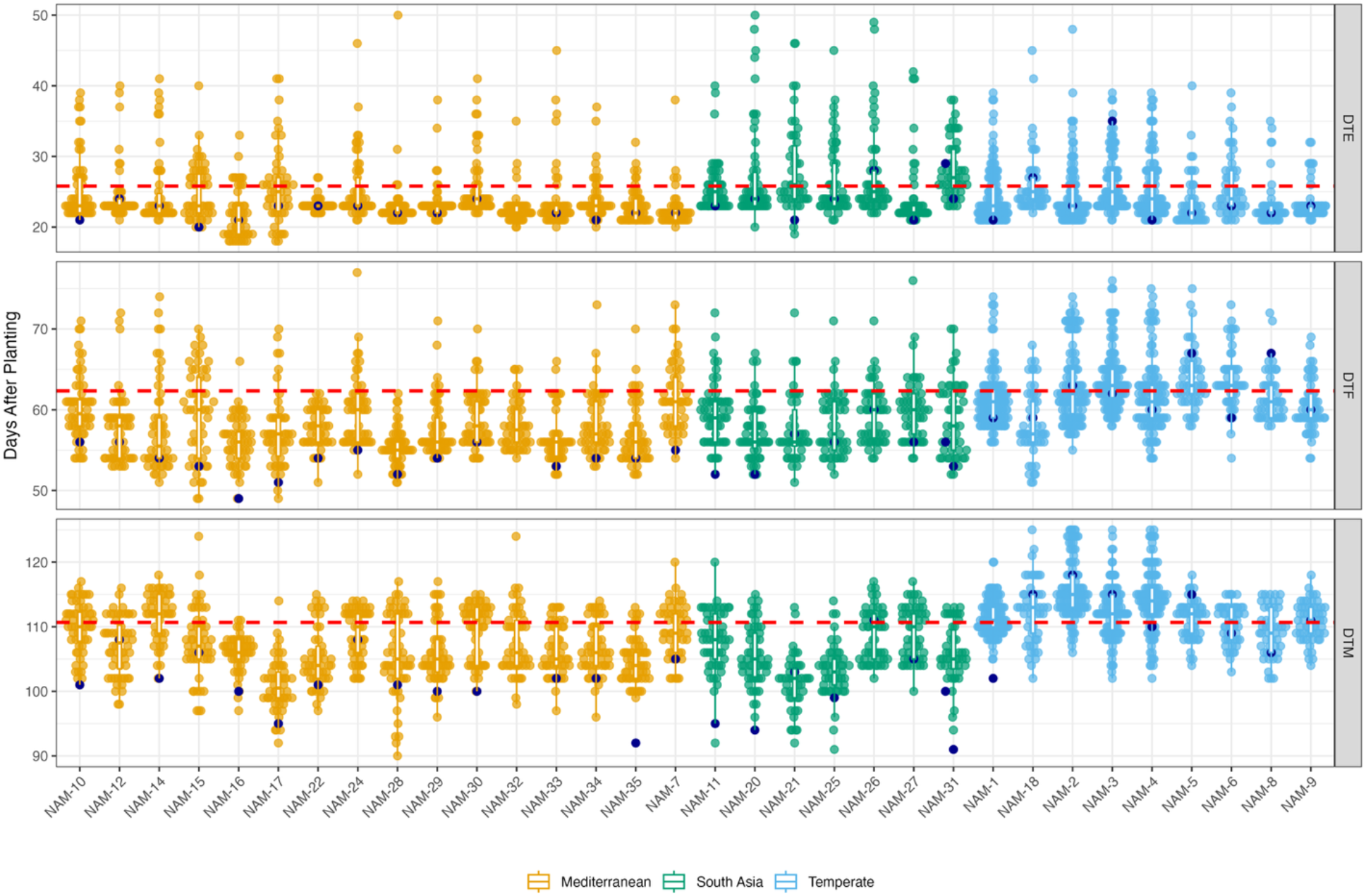
Variation in phenological stages, days to emergence (DTE), days to flowering (DTF), and days to maturity (DTM), among 32 RIL families, colored according to the macro-environmental origin of the parent crossed with CDC Redberry. The X-axis represents the RIL families, while the Y-axis indicates days after planting. Each dot represents an individual line within a specific RIL family, and the width of each facet shows the density distribution. The whiskers of the box plots represent 1.5 times the interquartile range. The red dashed horizontal line indicates the respective DTE, DTF and DTM for CDC Redberry (the common parent), while the dark blue dots highlight DTE, DTF and DTM of the second parent in each RIL family.

### 3.3 High-density genotyping resource for trait dissection

Genotyping of 2,017 lines (1,984 RILs and 33 parents) produced a high-quality set of 45,191 single nucleotide polymorphisms (SNPs), with an average marker density of 13 SNPs per megabase (Mb) across the CDC Redberry v2.0 reference genome (Ramsay et al., 2021). SNP distribution varied among chromosomes, ranging from 10 SNPs/Mb on chromosome 1 to 17 SNPs/Mb on chromosome 3 (Figure S2; Table S2).

This high-density SNP dataset represents a valuable genomic resource for lentil research, providing sufficient resolution for fine mapping of quantitative trait loci (QTL) and enabling genome-wide association studies (GWAS) with improved power and precision. The complete genotypic dataset is publicly available through the project page on our <u>KnowPulse</u> web portal, facilitating open-access use by researchers and breeders worldwide. By offering uniform, high-resolution genomic coverage, this dataset lays the foundation for dissecting complex traits, identifying candidate genes, and supporting marker-assisted and genomic selection strategies in lentil breeding programs.

### 3.4 More QTLs could be identified with the NAM

A Genome-wide association analysis (GWAS) using a high-quality SNP marker and for days to flowering (DTF as phenotypic data identified 14 significant marker-trait associations (MTAs)), located across chromosomes 1, 2, 3, 5, 6, and 7 (Figure 2; Table S3). As anticipated, the largest cluster of significant SNPs was detected on chromosome 6, near the *FTb* gene, a known regulator of flowering time in temperate environments (Haile et al., 2021; Neupane et al., 2022). The rediscovery of *FTb* validates the mapping approach and confirms the utility of the NAM population for detecting major-effect loci.

**Figure 2.**
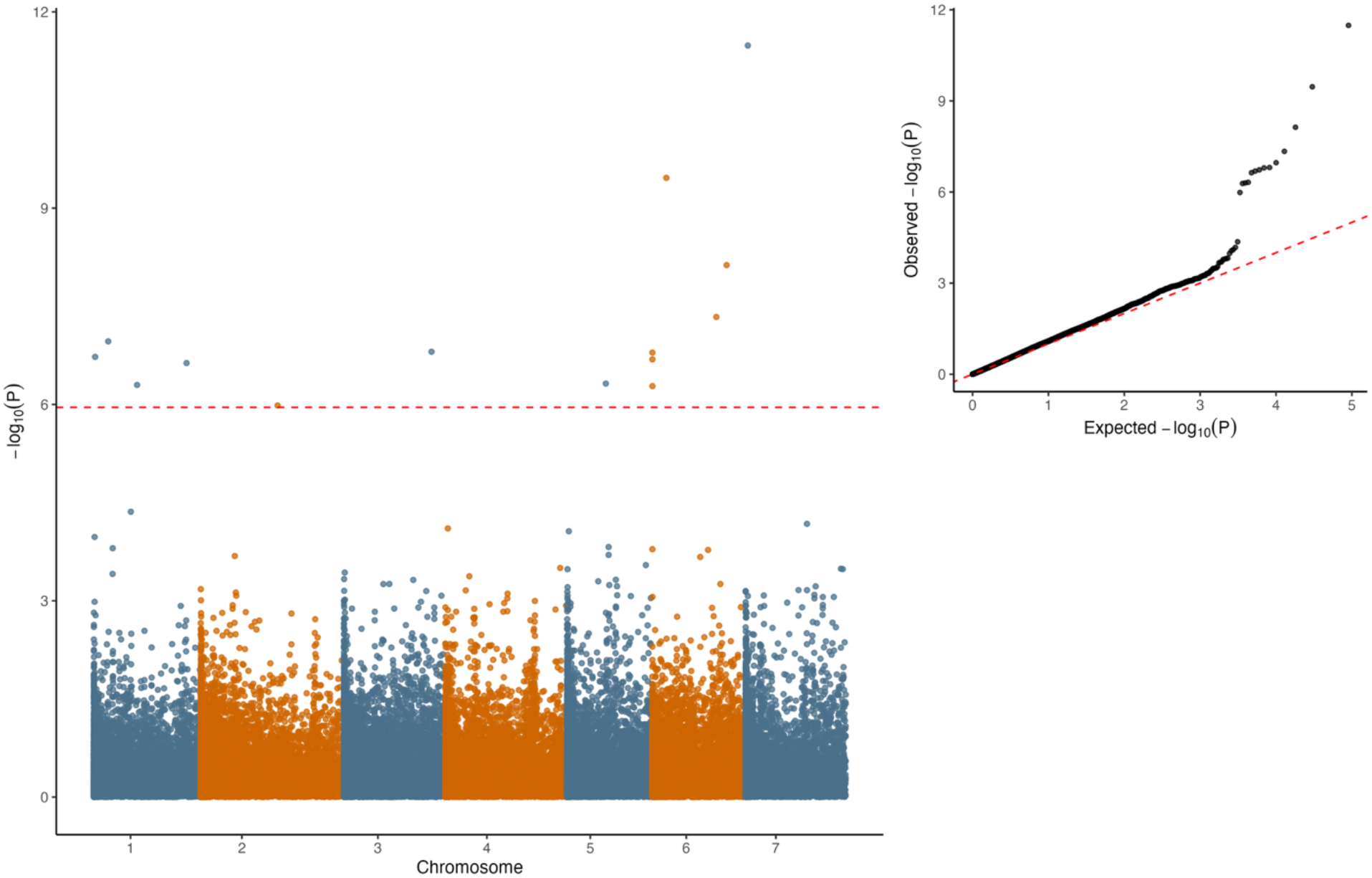
Genome-wide association analysis results for days to flowering (DTF) in the lentil NAM population, based on the 2019 field trial in Sutherland, SK. The left panel shows a Manhattan plot generated using the Bayesian-information and linkage disequilibrium iteratively nested keyway (BLINK) model. The horizontal red line indicates the significance threshold based on Bonferroni correction. The right panel presents a quantile–quantile (Q-Q) plot comparing observed vs. expected −log10(p) values. Significant right-tailed deviations in the Q-Q plot suggest robust marker-trait associations across the genome.

In addition to this known locus, a few novel QTLs were identified within 100 Kb of genes previously implicated in flowering regulation in other species, including:-

- Lcu.2RBY.Chr3_416260177 – located near *AP3a*, a MADS-box gene involved in floral organ development and early flowering transition, with homologous functions reported in *Glycine max* and *Medicago truncatula* (Roque et al., 2013; Zhang et al., 2023).
- Lcu.2RBY.Chr5_427555693, near *HUB2a*, which involved in epigenetic regulation of flowering via histone modification in *Arabidopsis thaliana* (Cao et al., 2008).
- Lcu.2RBY.Chr6_406916712, adjacent to *PIF6*, a gene influencing the flowering time by modulating photoperiod-dependent gene expression during floral transition in *Arabidopsis thaliana* (Howe, 2024)

These associations were not previously detected in diversity panel-based GWAS in lentil (Neupane et al., 2022), underscoring the enhanced resolution and detection power of the NAM design, particularly for identifying minor-effect loci and uncovering novel candidate regions.

## 4. Conclusion

This study reports the development of a genetically diverse and structurally informative lentil NAM population, designed as a resource for global lentil improvement. By integrating germplasm from diverse agro-ecological regions into a mappable framework, the population advances both genetic conservation and high-resolution trait dissection. The identification of both known loci (e.g., *FTb*) and novel candidate genes (e.g., *AP3a, HUB2a, PIF6*) for flowering time underscores the power of the NAM design in uncovering the genetic architecture of complex traits. Critically, the population structure enables the detection and pyramiding of minor-effect alleles, addressing limitations of traditional biparental QTL mapping and GWAS based on diversity panels. Although the present analysis was conducted in a single environment, future multi-year, multi-location trials integrated with multi-omics approaches are recommended to validate QTL stability and dissect genotype by environment interactions for flowering time and other complex traits. As multi-parent populations gain traction across crop species, this NAM population offers a timely and robust tool for dissecting complex traits and accelerating genetic gain in lentil improvement. Public availability of this resource will facilitate collaborative breeding, trait discovery, and pre-breeding efforts worldwide.

## Supporting information

https://knowpulse.usask.ca/research-study/AGILE-NAM-UAV-growth-modelling

## 5. Acknowledgements

This work was supported by the “Application of Genomics to Innovation in the Lentil Economy (AGILE),” project funded by Genome Canada and managed by Genome Prairie. We are grateful for the matching financial support from Western Grains Research Foundation, the Government of Saskatchewan, and the University of Saskatchewan. We acknowledge the technical assistance of the bioinformatics, field and molecular lab staff of the Pulse Crop Breeding and Genetics group at the University of Saskatchewan.

## 6. Author Contributions

S. Neupane: Data wrangling, methodology, writing original draft, writing review & editing.

R. Stonehouse: Genotyping, writing materials and methods of genotyping section.

L. Ramsay and M. Baker: Read mapping, SNP calling, quality analysis.

T. Haile: Methodology, designing and implementation of 2019 field trials.

D. J. Konkin and C. Sidebottom: Exome capture protocol for multiplexing large numbers of samples

K. E. Bett: Conceptualization, funding acquisition, methodology, writing review & editing, project administration, resource management, supervision.

## 7. Conflict of Interest

The authors declare no conflict of interest.

## 8. Data Availability

The data supporting this study are available at https://knowpulse.usask.ca/research-study/AGILE-NAM-UAV-growth-modelling or from the authors upon request.

## Notes

https://knowpulse.usask.ca/research-study/AGILE-NAM-UAV-growth-modelling

### Competing Interest Statement

The authors have declared no competing interest.

