## Supplementary material for "Development of a Nested Association Mapping (NAM) population for untangling complex traits in lentil (*Lens culinaris* Medik.)": https://knowpulse.usask.ca/research-study/AGILE-NAM-UAV-growth-modelling

**Supplementals:**

**Table 1.** Details of the generational stages of NAM used in genotyping and sequencing (N = 2047 plus 32 parents)

| Generation | Number of NAM Lines | Percentage (%) |
|---|---|---|
| F4:F5 | 2 | 0.10 |
| F5:F6 | 13 | 0.64 |
| F5:F7 | 2 | 0.10 |
| F6:F7 | 279 | 13.63 |
| F6:F8 | 27 | 1.32 |
| F7:F8 | 1473 | 71.96 |
| F7:F9 | 251 | 12.26 |
| Parents | 32 | |

**Table 2.** SNP count, chromosome length and SNP density of 45191 SNP markers distributed on seven chromosomes based on the CDC Redberry v2.0 genome (Ramsay et al., 2021).

| Chromosome | No. of SNPs | Length of assembled Chr (Mb) | SNP density (SNPs/Mb) |
|---|---|---|---|
| 1 | 5344 | 538.26 | 9.93 |
| 2 | 8116 | 613.63 | 13.23 |
| 3 | 7068 | 429.91 | 16.44 |
| 4 | 7119 | 482.06 | 14.77 |
| 5 | 6139 | 474.86 | 12.93 |
| 6 | 5707 | 420.42 | 13.57 |
| 7 | 5698 | 528.91 | 10.77 |

**Table 3.** Genome-wide association studies (GWAS) results

| SNP | Chr | Pos | P.value |
|---|---|---|---|
| Lcu.2RBY.Chr1_331862918 | 1 | 331862918 | 1.085636e-07 |
| Lcu.2RBY.Chr1_78037525 | 1 | 78037525 | 1.876298e-07 |
| Lcu.2RBY.Chr1_531219205 | 1 | 531219205 | 2.330866e-07 |
| Lcu.2RBY.Chr1_443206161 | 1 | 443206161 | 5.033340e-07 |
| Lcu.2RBY.Chr2_508328513 | 2 | 508328513 | 1.038197e-06 |
| Lcu.2RBY.Chr3_416260177 | 3 | 416260177 | 1.564089e-07 |
| Lcu.2RBY.Chr5_427555693 | 5 | 427555693 | 4.797130e-07 |
| Lcu.2RBY.Chr6_259785952 | 6 | 259785952 | 3.445885e-10 |
| Lcu.2RBY.Chr6_406916712 | 6 | 406916712 | 7.416910e-09 |
| Lcu.2RBY.Chr6_395972637 | 6 | 395972637 | 4.599805e-08 |
| Lcu.2RBY.Chr6_5134822 | 6 | 5134822 | 1.611981e-07 |
| Lcu.2RBY.Chr6_3607290 | 6 | 3607290 | 2.047135e-07 |
| Lcu.2RBY.Chr6_6335042 | 6 | 6335042 | 5.242370e-07 |
| Lcu.2RBY.Chr7_97102480 | 7 | 97102480 | 3.270548e-12 |

**Figures:**

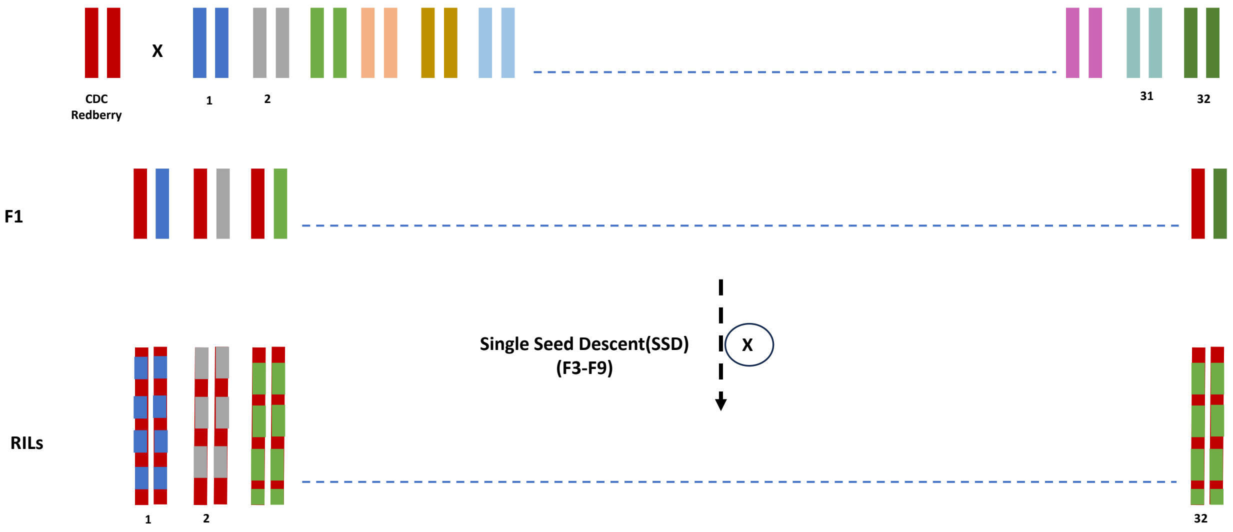


**Fig. 1.** Schematic representation of lentil NAM population development.

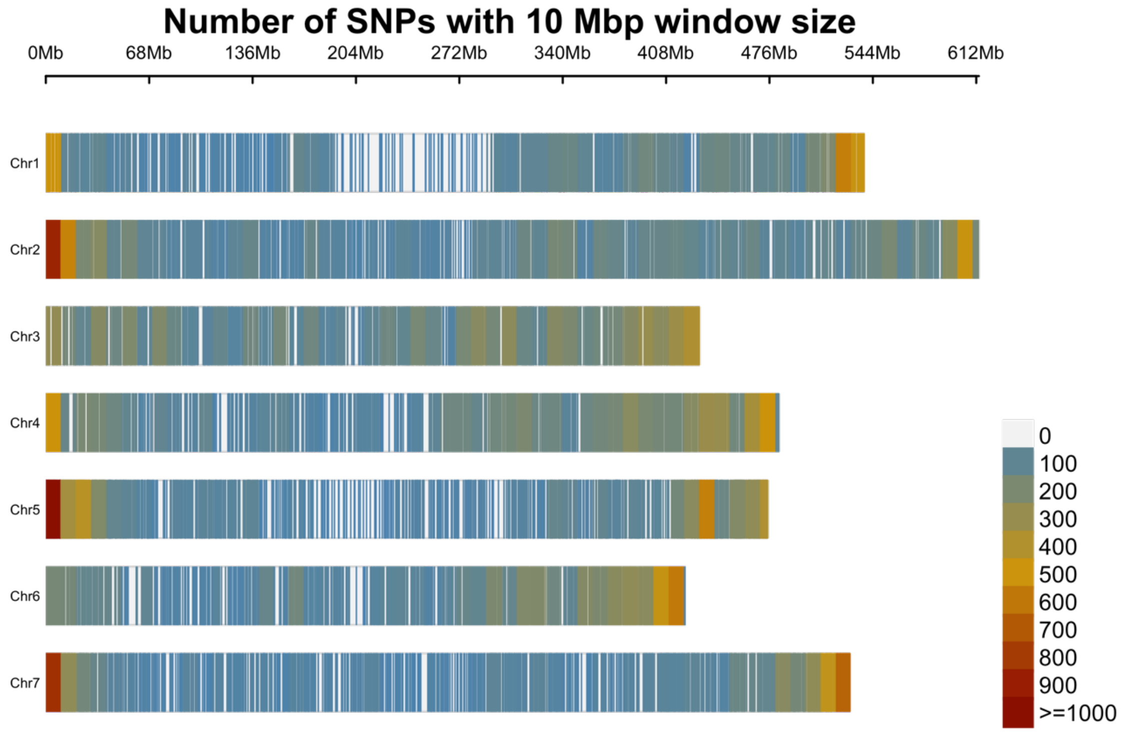


**Fig. 2.** Distribution of 45,191 single nucleotide polymorphisms (SNPs) across the seven chromosomes of the lentil on the CDC Redberry v2.0 genome (Ramsay et al., 2021). Colour represents marker density per 10-megabase pair window.
